# Intraneuronal chloride levels encode tiredness in cortex

**DOI:** 10.1101/2021.05.14.444189

**Authors:** Hannah Alfonsa, Paul Brodersen, Sarah E. Newey, Tomoko Yamagata, Marios C. Panayi, David M. Bannerman, Vladyslav V. Vyazovskiy, Colin J. Akerman

**Author notes:** Corresponding authors: Hannah Alfonsa, Colin J. Akerman.

## Abstract

Continuous periods of wakefulness are associated with reduced performance levels due to the build-up of sleep pressure in active regions of the brain. These effects manifest as use-dependent changes in cortical network activity and the mechanisms underlying these changes represent targets for overcoming the cognitive effects of tiredness. Here we reveal a central role for intraneuronal chloride levels, which increase in a use-dependent manner during waking, and reduce the strength of local synaptic inhibition in mouse cortex. Activity-dependent increases in chloride account for spatial and temporal features of sleep pressure, they underlie cortical network oscillations in the sleep-deprived state, and targeting chloride regulation in cortex can rescue performance levels when tired. These findings provide a missing link between sleep-wake history, synaptic transmission and cortical dynamics.

**One-Sentence Summary:** The effects of sleep pressure on cortical function are caused by use-dependent changes in chloride-mediated synaptic inhibition.

## Main Text

The increase in sleep pressure, or ‘need to sleep’, is associated with reduced cognitive performance and changes in cortical activity that can be detected first during the sleep-deprived awake state, and then during subsequent sleep (*1–4*). Whilst significant progress has been made in understanding the changes that accompany waking and sleeping (*5–8*), we still lack an appreciation of the cellular mechanisms that convert an animal’s sleep-wake history, into the changes in cortical activity and behaviour associated with tiredness. Understanding these mechanisms would reveal how the brain encodes the activity-dependent effects of wakefulness and reversing such mechanisms offers the potential to reset performance levels when we are tired. The effects of wakefulness vary temporally and spatially across cortex, in a manner that reflects the duration and task-dependent demands of the preceding wake period (*2, 4, 9, 10*). This is evident in the level of slow wave activity (SWA) recorded in the cortical electroencephalogram (EEG; spectral power between 0.5 to 4 Hz) during non-rapid eye movement (NREM) sleep, which represents the most reliable marker of sleep pressure (*4, 9*). Not only does SWA intensify as a function of preceding wakefulness (*9*), but it exhibits local regulation such that regions most active during the preceding wake period, show the greatest increases in SWA (*10–14*). In a related manner, wakefulness leads to low frequency cortical oscillations in the sleep-deprived state (*1, 15, 16*). Referred to as ‘local sleep’ in the awake brain, this phenomenon is also associated with task-dependent demands and has been proposed to compromise cortical function when we are tired (*1, 3, 17*). Here we reveal that the regulation of cortical intraneuronal chloride concentration ([Cl^-^]_i_) plays a central role in the underlying mechanisms. [Cl^-^] determines the strength of fast inhibitory synaptic transmission mediated by Cl^-^-permeable γ-aminobutyric acid type A (GABA_A_) receptors (GABA_A_Rs). We find that local, activity-dependent increases in cortical [Cl^-^]i during wakefulness determine the amount of SWA during NREM sleep, the intensity of low frequency oscillations in the sleep-deprived state, and levels of behavioural performance when tired.

### Waking is associated with activity-dependent increases in cortical [Cl^-^]_i_

We performed experiments in mice to assess cortical [Cl^-^]_i_ at different time points during the 24-hour light-dark cycle. These time points were associated with different sleep-wake histories, as confirmed by continuous EEG and EMG recordings (Fig. 1A). The mice spent a high proportion of time asleep during the light period, whereas they spent a high proportion of time awake during the dark period, and this period of wakefulness could be extended by performing a sleep deprivation (SD) protocol at the beginning of the light period (Fig. 1A; see Methods). [Cl^-^]i measurements were performed in acute somatosensory cortex (S1) brain slices prepared from (i) mice 3 hours after light onset (i.e. Zeitgeber time 3, ZT3), which is associated with recent sleep, (ii) mice 3 hours after dark onset (ZT15), which is associated with recent waking, or (iii) after 3 hours of SD that started at light onset (ZT3-SD). The ZT3-SD condition served as a control for potential circadian processes, as it shares the same ZT as the ZT3 condition, but has a different sleep-wake history.

**Fig. 1.**
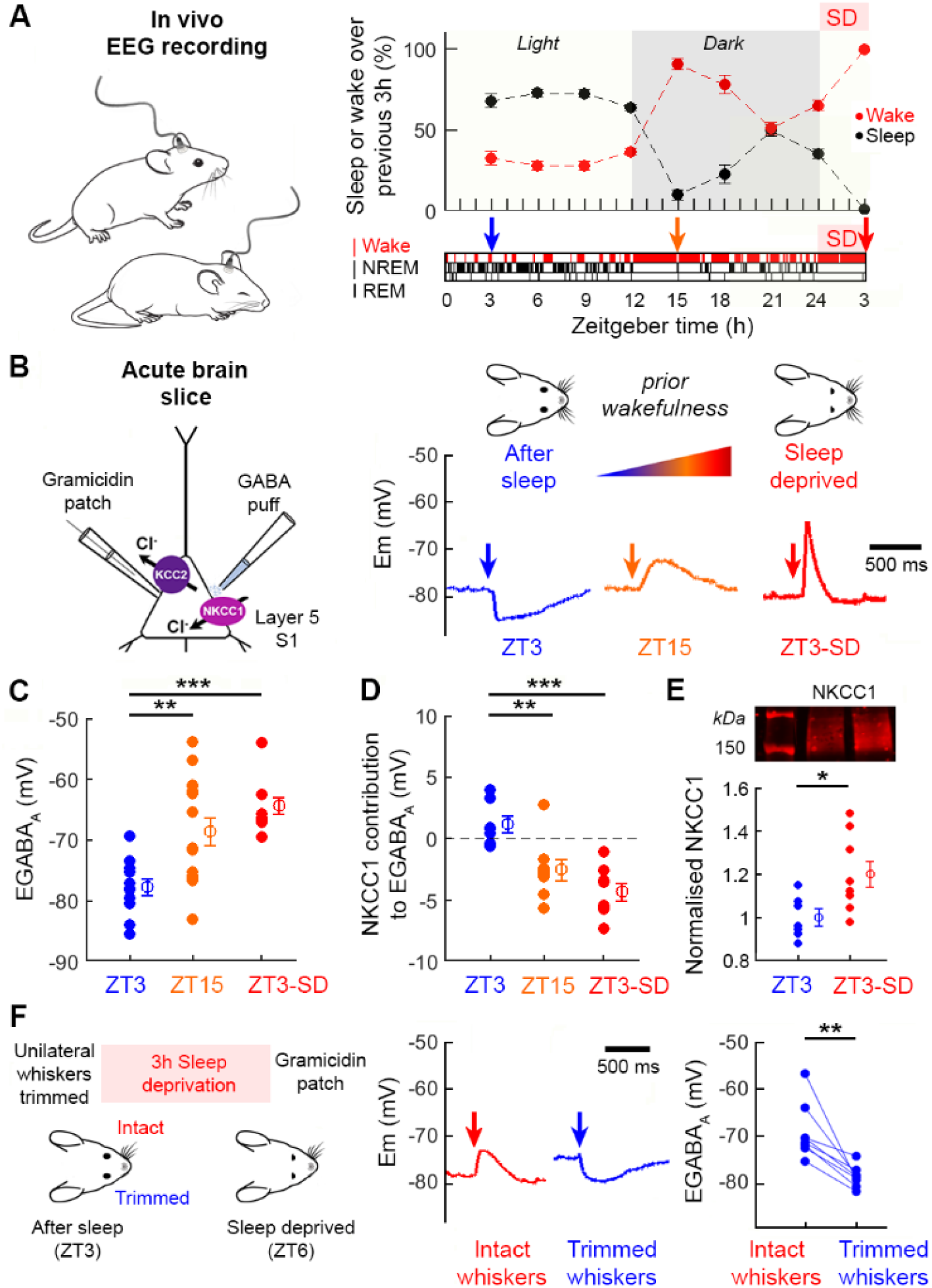
Waking is associated with activity-dependent increases in cortical [Cl^-^]_i_. (**A**) Continuous EEG recordings were performed in freely moving mice (left). Population data (right) shows average time spent asleep or awake over 3-hour intervals (6 animals, 10 days) either without, or with, a period of sleep deprivation (SD; 5 animals, 5 days) at light onset. Hypnogram shows typical distribution of wake, NREM and REM sleep. Arrows indicate when cortical [Cl^-^]_i_ was measured. (**B**) Gramicidin perforated patch recordings compared GABAAR signalling in L5 pyramidal neurons from acute slices of primary somatosensory cortex (S1; left). Example current clamp recordings (right) show the effect of GABA_A_R activation at ZT3, ZT15, and ZT3 after a 3-hour SD protocol (‘ZT3-SD’). (**C**) EGABA_A_ was more depolarized at ZT15 (3 animals, 15 neurons; **p<0.005, unpaired t-test) and ZT3-SD (4 animals, 10 neurons; ***p<0.0001, unpaired t-test) compared to ZT3 (3 animals, 11 neurons). (**D**) Blocking NKCC1 with bumetanide induced larger EGABA_A_ shifts at ZT15 (4 animals, 8 neurons; **p<0.003, unpaired t-test) and ZT3-SD (3 animals, 8 neurons; ***p<0.0001, unpaired t-test) compared to ZT3 (5 animals, 8 neurons). (**E**) Representative western blot (top) comparing NKCC1 levels in S1 of ZT3 and ZT3-SD mice. Normalized NKCC1 levels (bottom) increased in the ZT3-SD condition (7 and 8 animals, 3 blots; *p<0.02, unpaired t-test). (**F**) Sensory deprivation paradigm involved trimming whiskers unilaterally at ZT3, when [Cl^-^]_i_ is normally low, followed by 3 hours of SD (left). Example current clamp recordings from the same animal (middle) comparing GABAAR activation in S1 receiving intact whisker input and the contralateral S1, with trimmed whisker input. EGABA_A_ (right) was more hyperpolarized in the hemisphere deprived of whisker input (3 animals, 8 paired neurons; **p<0.005, Wilcoxon test).

To measure [Cl^-^]_i_, layer 5 (L5) pyramidal neurons were targeted using the gramicidin perforated patch recording technique, which avoids disruption of transmembrane Cl^-^ gradients so that GABA_A_R activity reflects the neuron’s native [Cl^-^]_i_ (Fig. 1B). Recordings in current clamp mode revealed that neurons from slices prepared at ZT3 showed more hyperpolarizing GABA_A_R responses to brief puffs of GABA, whilst neurons from slices prepared at ZT15 and ZT3-SD exhibited more depolarizing GABA_A_R responses (Fig. 1B). No difference was observed in resting membrane potential across the conditions (ZT3: −77.0 ± 2.1 mV, ZT15: −76.5 ± 1.8 mV and ZT3-SD: −75.6 ± 3.5 mV). These observations were confirmed by measurements of the GABA_A_R reversal potential (EGABA_A_) in voltage clamp mode, which directly reflects [Cl^-^]_i_ (Fig. S1). Mice at ZT3 exhibited hyperpolarized EGABA_A_ values, reflecting low [Cl^-^]_i_. Whereas mice at ZT15 and ZT3-SD exhibited more depolarized EGABA_A_, reflecting higher [Cl^-^]_i_ (Fig. 1C). The fact that EGABA_A_ at ZT3-SD was significantly more depolarized than ZT3, established that the change in cortical [Cl^-^]_i_ reflects sleep-wake history and not circadian processes. Furthermore, equivalent recordings from L2/3 pyramidal neurons in auditory cortex showed comparable differences, revealing that the relationship between sleep-wake history and [Cl^-^]i was not unique to cortical layer or region (Fig. S2). Whilst control experiments confirmed that the observed differences were independent of spiking activity and the endogenous activation of GABA_A_Rs within the slices at the time of recording (Fig. S2). Therefore, [Cl^-^]i in mouse cortex reflects the recent sleep-wake history of the animal, with wakefulness being associated with increases in [Cl^-^]i and more depolarized EGABA_A_.

[Cl^-^]_i_ is determined by cotransporter proteins in the cell membrane and the relative activity of oppositely directed cotransporters can generate temporal and spatial differences in a neuron’s [Cl^-^]_i_ (*18*). To investigate the mechanism underlying sleep-wake related cortical [Cl^-^]_i_ dynamics, we measured the contribution of the potassium-chloride exporter, KCC2, and the sodium-potassium-chloride importer, NKCC1, by blocking each cotransporter with either VU0463271 (‘VU’) or bumetanide, respectively. Whereas the KCC2 contribution to EGABA_A_ remained stable (Fig. S3), the contribution of NKCC1 to EGABA_A_ changed depending on the sleep-wake history (Fig. 1D). Blocking NKCC1 with bumetanide had no effect upon the hyperpolarized EGABA_A_ values at ZT3, but bumetanide caused a significant hyperpolarizing shift in EGABA_A_ when applied at ZT15 or ZT3-SD, consistent with a greater contribution of NKCC1 to EGABA_A_ after periods of wakefulness (Fig. 1D). Western blots supported these observations by revealing an increase in NKCC1 protein levels in S1 at ZT3-SD compared to ZT3 (Fig. 1E).

Finally, to assess whether the wakefulness-associated increases in [Cl^-^]_i_ are use-dependent, we used a sensory deprivation paradigm that involved trimming whiskers unilaterally at ZT3, when [Cl^-^]_i_ is normally low. Mice were then subjected to 3 hours of SD so that they would remain awake and able to use whiskers on the intact side, but not on the trimmed side (Fig. 1F). This protocol resulted in a shift towards depolarizing GABA_A_R responses in S1 L5 pyramidal neurons recorded in the hemisphere receiving intact whisker input. Whereas S1 neurons recorded in the same brain slices, but from the hemisphere deprived of whisker input, continued to exhibit hyperpolarizing GABA_A_R responses, consistent with a low [Cl^-^]_i_ (Fig. 1F). In conclusion therefore, wakefulness is associated with use-dependent increases in cortical [Cl^-^]_i_.

### Cortical [Cl^-^]_i_ determines the level of local NREM SWA

Given the central role for synaptic inhibition in regulating network activity, we wondered whether the observed changes in cortical [Cl^-^]_i_ regulate the level of NREM SWA, which is the most reliable marker of sleep pressure and reflects both the duration and task-dependent demands of preceding wake periods (*2, 4, 9, 10*). If wakefulness-associated increases in [Cl^-^]_i_ regulate the level of cortical SWA, our prediction was that lowering local [Cl^-^]_i_ in a region of cortex would have its greatest impact when sleep pressure is high, at the beginning of light onset. To test this, we performed *in vivo* recordings in freely moving mice that had been implanted with an LFP electrode coupled with an infusion cannula, and targeted to L5 of S1 (Fig. 2A). Mice were maintained on a standard 24-hour light-dark cycle and continuous LFP recordings were performed to calculate LFP spectral power during NREM sleep. Antagonists of either NKCC1 or KCC2 were then infused locally into S1 at different time points during the light period, which are associated with different levels of SWA (Fig. 2B). Lowering [Cl^-^]_i_ by blocking NKCC1 with bumetanide at the beginning of the light period, when SWA is highest, caused pronounced reductions in SWA levels in S1 (Fig. 2C-D). These effects were only evident in the local S1 LFP signal and not in the global frontal EEG (Fig. 2D), consistent with the idea that [Cl^-^]_i_ can mediate regional differences in SWA. In contrast, 2-3 hours later, when SWA levels have reduced as a result of sleep following the onset of light, the same manipulation did not affect local or global SWA levels (Fig. 2E-F). Increasing [Cl^-^]_i_ at this later time point by blocking KCC2 with VU, was sufficient to cause an increase in the level of local SWA without affecting global EEG (Fig. 2G-H). Furthermore, by using the same sensory deprivation paradigm as described above, we confirmed previous reports (*10*) that high NREM SWA levels reflect activity-dependent processes during waking (Fig. S4). Again, locally raising [Cl^-^]_i_ in this context with VU, was sufficient to rescue the high SWA levels that were prevented by sensory deprivation (Fig. S4). Taken together, these data support the conclusion that cortical [Cl^-^]_i_ regulates NREM SWA levels.

**Fig. 2.**
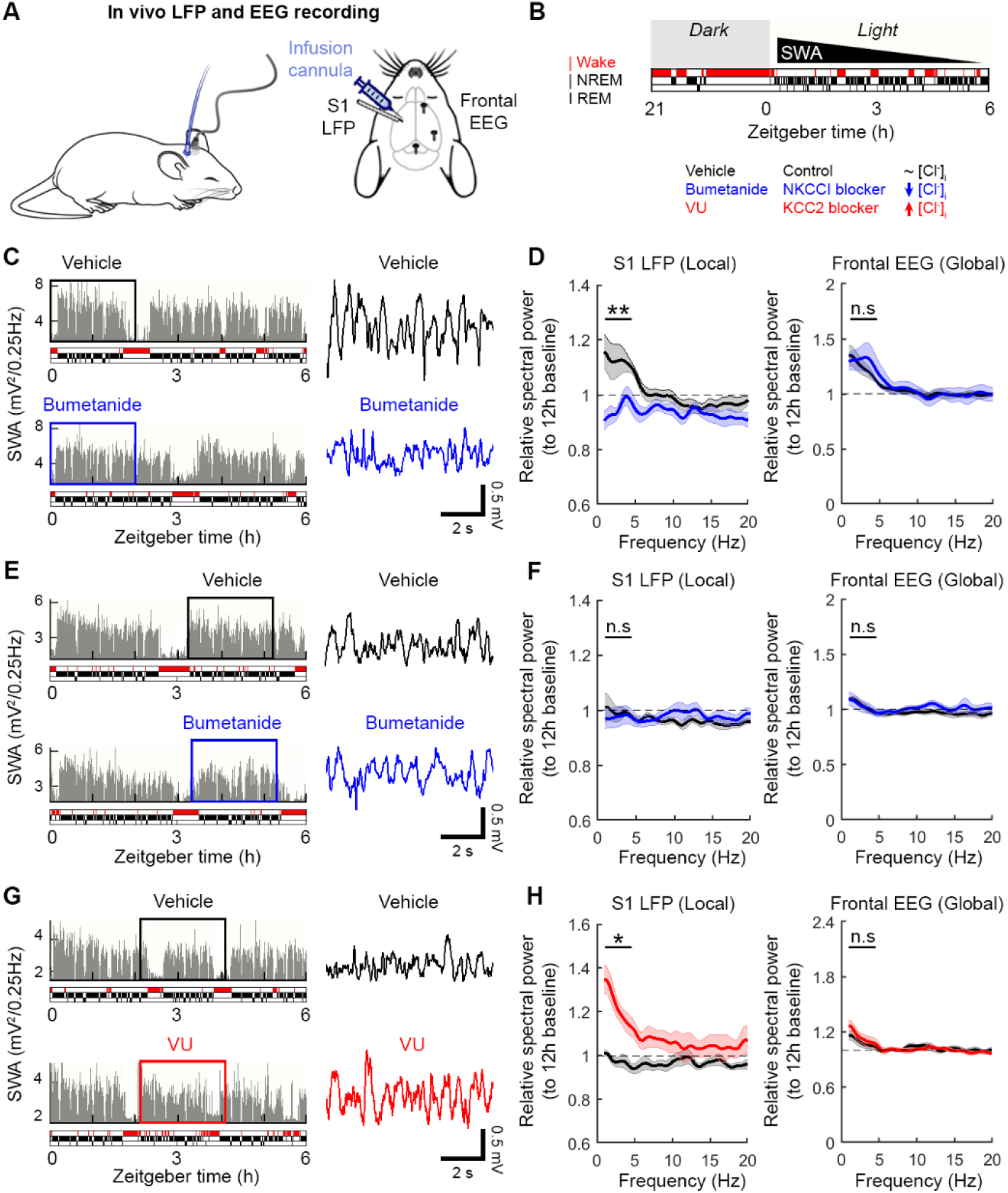
Cortical [Cl^-^]_i_ determines the level of local NREM SWA. (**A**) An LFP electrode coupled to an infusion cannula was targeted to L5 of left S1 and frontal EEG screws were positioned over the right hemisphere. Continuous LFP and EEG recordings were used to monitor local and global spectral power, respectively. (**B**) NREM SWA (black triangle) is most intense at sleep onset (coincident with light onset) and then lessens over the course of sleep. [Cl^-^]_i_ was manipulated at different points during NREM sleep, by locally infusing blockers of NKCC1 or KCC2. (**C**) LFP in a mouse that received vehicle (top) or bumetanide (bottom) during early NREM sleep, on different days. Expanded traces (right) are from the boxed regions. (**D**) Bumetanide infusion during early NREM sleep reduced local spectral power in the SWA range (left; 5 animals, 6 trials; **p<0.003, paired t-test), without affecting frontal EEG (right; p=0.39, paired t-test). (**E**) LFP in a mouse that received vehicle (top) or bumetanide (bottom) during later NREM sleep. (**F**) Bumetanide infusion during later NREM sleep did not affect local spectral power in the SWA range (left; 5 animals, 6 trials; p=0.8, paired t-test), or frontal EEG (right; p=0.97, paired t-test). (**G**) LFP in a mouse that received infusion of vehicle (top) or VU (bottom) during later NREM sleep. (**H**) VU infusion during later NREM sleep increased local spectral power in the SWA range (left; 4 animals, 6 trials; *p<0.01, paired t-test), without affecting frontal EEG (right; p=0.35, paired t-test).

### High [Cl^-^]_i_ weakens local synaptic inhibition and enhances neuronal recruitment during SWA

SWA involves the low frequency switching between periods of action potential firing (ON periods, associated with cortical up states) and periods of neuronal silence (OFF periods, associated with cortical down states) (*9, 19*), with higher SWA levels reflecting greater spiking activity and more synchronous recruitment during ON periods (*9*). As ON periods involve the dynamic interplay between inhibitory and excitatory neurotransmission (*20–22*), we postulated that neurons with elevated [Cl^-^]_i_ would experience weaker synaptic inhibition and therefore more readily summate excitatory input to reach spike threshold (*23*). To test this, we generated brain slices from animals that had experienced different sleep-wake histories (ZT3 versus ZT15) and used cell-attached recordings to measure the spike probability of L5 pyramidal neurons following stimulation of two independent input pathways (Fig. 3A; Fig. S5). Neurons at ZT15 with more depolarized EGABA_A_ showed broader synaptic integration windows than neurons at ZT3, consistent with weaker synaptic inhibition in animals that had experienced recent wakefulness. Reducing [Cl^-^]_i_ in ZT15 neurons (by blocking NKCC1 with bumetanide) narrowed the synaptic integration window, whereas increasing [Cl^-^]_i_ in ZT3 neurons (by blocking KCC2 with VU) broadened the synaptic integration window (Fig. 3A). We also generated a simple network model, which enabled us to selectively vary [Cl^-^]_i_ in a population of excitatory neurons, whilst maintaining all other parameters (Fig. 3B; see Methods). Networks of neurons with more depolarized EGABA_A_ (i.e. higher [Cl^-^]_i_) showed greater recruitment of spiking activity during simulated ON periods, which was most pronounced when ON-OFF transitions occurred at lower frequencies (≤ 6 Hz; Fig. 3B). These data support the conclusion that, through their effects upon synaptic inhibition, [Cl^-^]_i_ levels associated with preceding wakefulness determine how easily a cortical neuron is recruited by excitatory synaptic input, as occurs during ON periods.

**Fig. 3.**
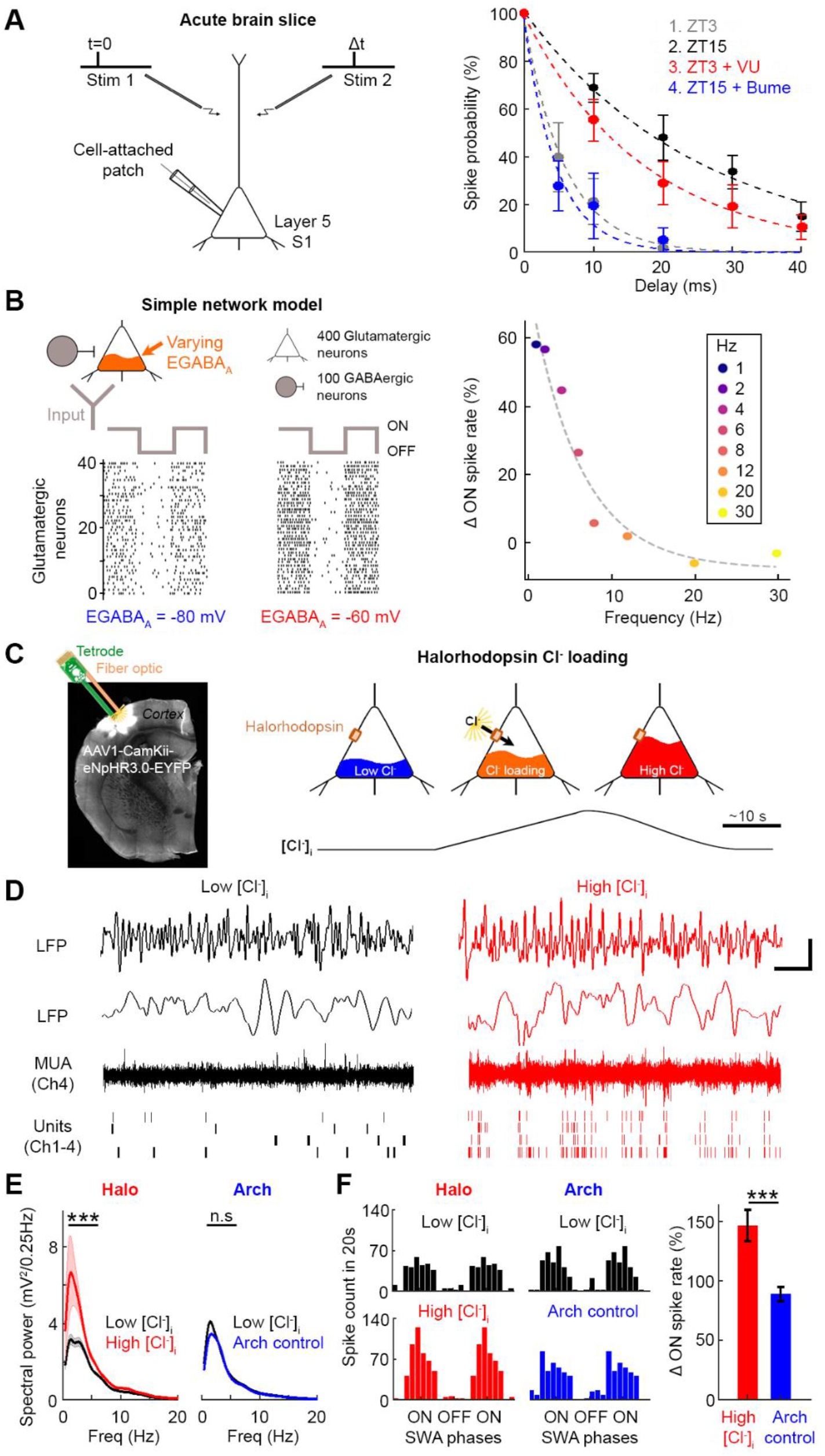
High [Cl^-^]_i_ weakens local synaptic inhibition and enhances neuronal recruitment during SWA. (**A**) L5 synaptic integration windows were measured from spiking responses to electrical stimulation of two input pathways (left). Normalized spike probability plotted for different inter-stimulus delays and conditions (right). Narrow integration windows were associated with ZT3 (grey: 3 animals, 8 neurons) and following NKCC1 blockade at ZT15 (blue: 3 animals, 6 neurons). Wider integration windows were associated with ZT15 (black: 3 animals, 10 neurons) and following KCC2 blockade at ZT3 (red: 4 animals, 8 neurons; multi-group comparison p<0.0001, one-way ANOVA; ZT3 vs. ZT15, ZT15 vs. ZT15 + Bume, and ZT3 vs. ZT3 + VU, all p<0.05, Bonferroni post-tests). (**B**) Simple network model comprising 400 glutamatergic and 100 GABAergic neurons, receiving synchronous oscillatory input to simulate ON and OFF periods. Raster plots show excitatory neuron spiking in response to a 1 Hz input (left). An EGABAA shift from −80 to −60 mV increased spiking during ON periods, especially at lower input frequencies (right). (**C**) Halorhodopsin (‘Halo’, eNpHR3.0) was expressed in S1 excitatory cortical neurons and a combined tetrode and fibre optic implant were targeted to L5 *in vivo* (left). Recordings were compared during the 20 s before light activation, when [Cl^-^]_i_ is ‘low’, then immediately after a 20 s period of Halo activation, when [Cl^-^]_i_ would be ‘high’ (right). (**D**) LFP (top) and multi-unit spiking (bottom) under low [Cl^-^]_i_ (left) and high [Cl^-^]_i_ (right) conditions. Scale bar, 2 s and 0.5 mV. (**E**) LFP spectral power in the SWA range increased after Halo activation (3 animals, 9 days, 35 trials; ***p<0.0001, Wilcoxon test), but not after activation of archaerhodopsin (‘Arch’), which served as a hyperpolarization control (2 animals, 5 days, 19 trials; p=0.25, paired t-test). (**F**) Phase-plots (left) of multi-unit spiking relative to SWA before (top) and after (bottom) activation of Halo or Arch. Halo-mediated Cl^-^ loading increased spike rate during ON periods of SWA (right; red: 25 trials; blue: 19 trials; ***p<0.001, unpaired t-test).

To test this mechanism in the intact brain, we used an *in vivo* optogenetic strategy to control [Cl^-^]_i_ in a population of cortical neurons, whilst assessing local SWA levels (Fig. 3C). Previous work has established that the light-activated Cl^-^ pump, halorhodopsin (eNpHR3.0), can be used to generate transient and physiologically-relevant increases in [Cl^-^]_i_ (*24, 25*). Virally-delivered halorhodopsin was expressed in pyramidal neurons of S1 and a tetrode coupled with fibre optic was implanted to monitor and manipulate nearby neurons. [Cl^-^]_i_ was increased optically during periods of NREM sleep at ZT3 when [Cl^-^]_i_ is normally low. LFP and multiunit activity were compared immediately before and after halorhodopsin activation, when levels of [Cl^-^]_i_ in the opsin-expressing neurons would be low and high, respectively (Fig. 3C). A high [Cl^-^]_i_ was associated with increased LFP spectral power in the lower frequency range (0.5 to 6 Hz; Fig. 3D-E), which recovered with a mean time constant of 12.7 ±1.5 s (Fig. S6), consistent with previous observations (*24, 25*). These effects were not observed with the light-activated outward proton pump, archaerhodopsin, which provided a control for membrane potential hyperpolarization without affecting [Cl^-^]_i_ (Fig. 3E and Fig. S6). To assess neuronal recruitment into population ON and OFF periods, the timing of multiunit activity relative to slow waves was analysed before and after opsin activation. Phase plots of spike-time histograms showed that halorhodopsin-mediated increases in [Cl^-^]_i_ produced an increase in spiking activity during ON periods, which was not evident following archaerhodopsin activation (Fig. 3F and Fig. S6). Taken together, elevated [Cl^-^]_i_ in a population of excitatory cortical neurons is sufficient to increase their synaptic recruitment during ON periods and boost local cortical SWA. [Cl^-^]_i_ therefore provides a direct mechanistic link between previous sleep-wake history and the spatial and temporal regulation of cortical SWA, which reflects sleep pressure.

### Wake-related increases in [Cl^-^]_i_ change low frequency cortical oscillations and compromise performance in the awake sleep-deprived state

Like humans, mice become sleepy with extended periods of wakefulness and show reduced cognitive and behavioural performance (*26, 27*). These changes have been linked to increases in low frequency cortical oscillations in the sleep-deprived awake EEG, which show spectral overlap with SWA, vary in a manner that reflects task-dependent demands, and have been seen as evidence of local sleep in the awake brain (*1, 3, 15–17*). To examine whether cortical [Cl^-^]_i_ dynamics underlie these phenomena, we first established that the awake S1 LFP and frontal EEG both show increased power at low frequencies (2 to 6 Hz) during a period of SD at the beginning of the light period Fig. 4A-C). If elevated [Cl^-^]_i_ promotes this activity in the sleep-deprived awake state, we predicted that manipulating Cl^-^ cotransporters in S1 would affect these signals locally, but not globally. In line with this prediction, infusion of bumetanide into S1 prevented the local increase in low frequency oscillatory activity during SD, but not in the frontal EEG (Fig. 4C). Whereas raising [Cl^-^]_i_ with local VU augmented low frequency activity in the S1 LFP, but not in the frontal EEG (Fig. 4C). To confirm the use-dependent nature of [Cl^-^]_i_ changes in this process, we used the same sensory deprivation paradigm as described above, and performed LFP/EEG measurements in the awake sleep-deprived state (Fig. 4D). Unilateral whisker trimming combined with SD prevented the build-up of awake low frequency oscillations in S1 of the sensory deprived hemisphere, but not globally or in S1 of the hemisphere that received intact sensory input during the period of SD (Fig. 4E). Furthermore, the effects of sensory deprivation could be rescued by raising [Cl^-^]_i_ via local infusion of VU into S1 of the sensory deprived hemisphere (Fig. 4E). Thus, activity-dependent increases in [Cl^-^]_i_ boost levels of low frequency oscillatory cortical activity in the sleep-deprived awake state, which reflect use-dependent demands during the preceding wake period.

**Fig. 4.**
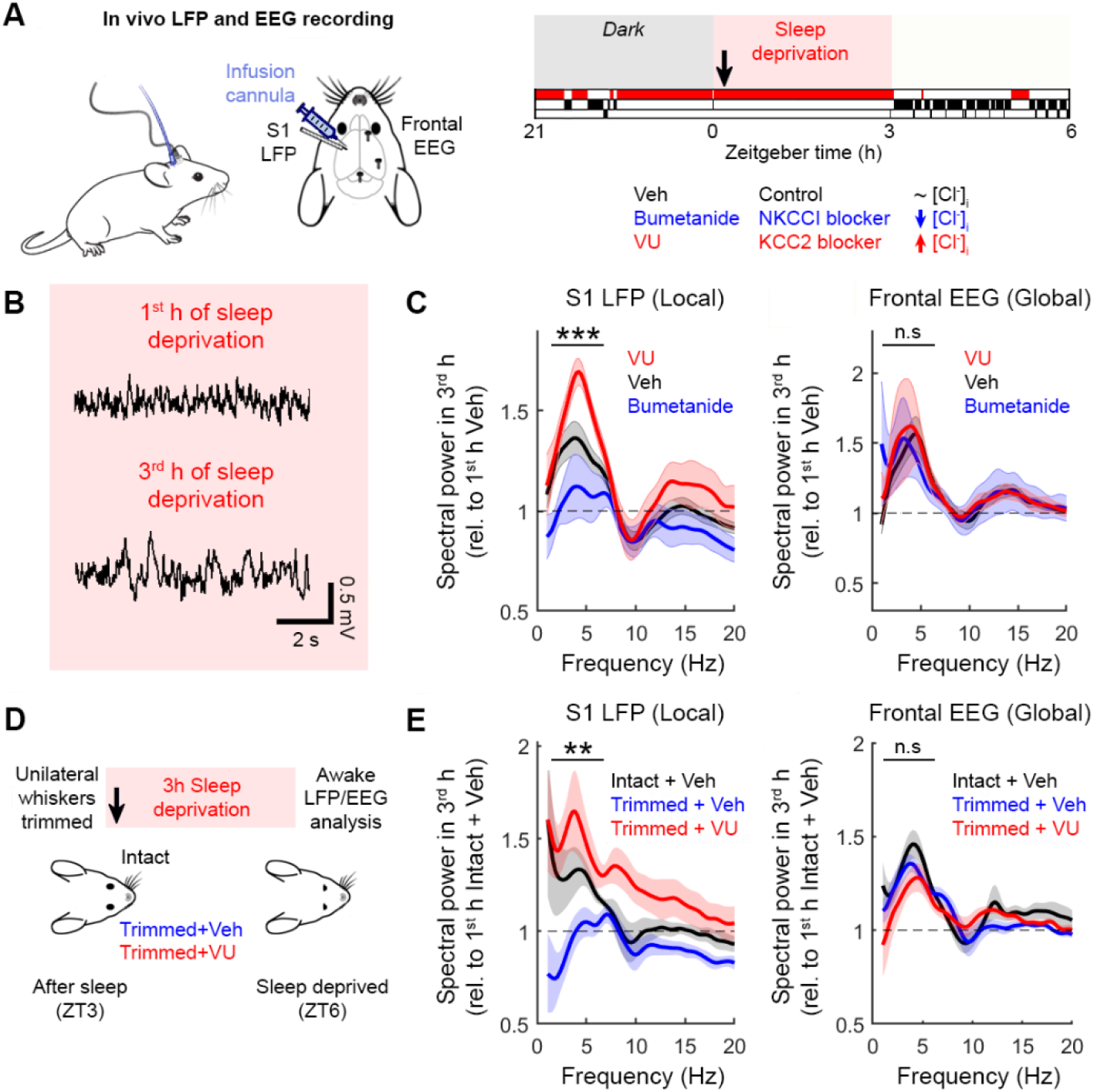
High [Cl^-^]_i_ increases low frequency cortical oscillations in the awake sleep-deprived state. (**A**) An LFP electrode coupled to an infusion cannula was targeted to L5 of left S1 and frontal EEG screws were positioned over the right hemisphere (left). Continuous awake LFP and EEG recordings were used to monitor local and global spectral power, respectively. Mice experienced a 3-hour SD protocol at the beginning of the light period (right), during which [Cl^-^]_i_ was manipulated by locally infusing blockers of NKCC1 or KCC2 (arrow indicates start of infusion). (**B**) Awake LFP traces from a control mouse show increase in low frequency cortical oscillations between the 1^st^ hour (top) and 3^rd^ hour (bottom) of SD. (**C**) Awake LFP (left) revealed an increase in low frequency cortical oscillations (2-6Hz; black: 15 animals), which was reduced by bumetanide (blue: 7 animals) and increased by VU (red: 8 animals; multi-group comparison ***p<0.001, one-way ANOVA; Veh vs. Bumetanide and Veh vs. VU, both p<0.005, paired t-test), without affecting frontal EEG (right; black: 13 animals, blue: 7 animals, red: 6 animals; multi-group comparison p=0.98, one-way ANOVA; Veh vs. Bumetanide, Veh vs. VU, p=0.76 and p=0.93, paired t-test). (**D**) Sensory deprivation paradigm combined unilateral whisker trimming at ZT3, followed by 3 hours of SD, during which either vehicle or VU was infused unilaterally. (**E**) Awake LFP (left) revealed that whisker trimming prevented the increase in local low frequency cortical oscillations (black: 8 animals, blue: 7 animals), which could be rescued by VU infusion into S1 (red: 7 animals; multi-group comparison **p<0.003, one-way ANOVA; Intact+Veh vs. Trimmed+Veh, p<0.01 paired t-test; Trimmed+Veh vs. Trimmed+VU, p<0.005, unpaired t-test). Neither whisker trimming nor S1 infusion affected the increase in low frequency oscillations detected in frontal EEG (right; black: 10 animals, blue: 7 animals, red: 8 animals; multi-group comparison p=0.13, one-way ANOVA; Intact+Veh vs. Trimmed+Veh, p=0.16 paired t-test; Trimmed+Veh vs. Trimmed+VU, p=0.36, unpaired t-test).

Given the role for [Cl^-^]_i_ in converting preceding wakefulness into the changes in cortical activity associated with high sleep pressure, an exciting possibility is that altering [Cl^-^]_i_ could reverse the drops in cognitive performance levels that are associated with tiredness (*1, 3, 17*). To test this, we investigated the impact of cortical [Cl^-^]_i_ upon performance levels on novel object recognition (NOR) tasks in sleep-deprived animals that had experienced 3 hours of SD at the beginning of light onset (Fig. 5A). Performance was compared to rested animals that were allowed to sleep during the equivalent 3-hour period and manipulations of [Cl^-^]_i_ were targeted to S1. By comparing performance across two different sensory modalities, we could also confirm whether effects were due to altering local cortical [Cl^-^]i, rather than a non-specific global effect. A tactile NOR task conducted in the dark was used to assess somatosensory cortical processing (Fig. 5B), whilst an odor-based NOR task served as a control for sensory modality that was remote from the infusion site (see Methods). First, we confirmed that SD resulted in decreased performance levels (*26, 27*), by demonstrating that SD resulted in lower novelty preference on both the tactile and odor NOR tasks (Fig. 5C, 5F and Fig. S7). If elevated [Cl^-^]_i_ and the associated increase in low frequency cortical oscillations contribute to these decreased performance levels, reducing [Cl^-^]_i_ in S1 would be expected to improve performance on the tactile task, but not on the odor task. Consistent with this prediction, blocking NKCC1 by local infusion of bumetanide into S1 of SD animals resulted in increased NOR performance on the tactile task, but not on the odor task (Fig. 5D and 5G). Finally, an increase in [Cl^-^]_i_ was also shown to be sufficient to induce the drops in performance level, as local VU infusion into S1 caused rested animals to show reduced NOR performance on the tactile task, but not on the odor task (Fig. 5E and 5H).

**Fig. 5.**
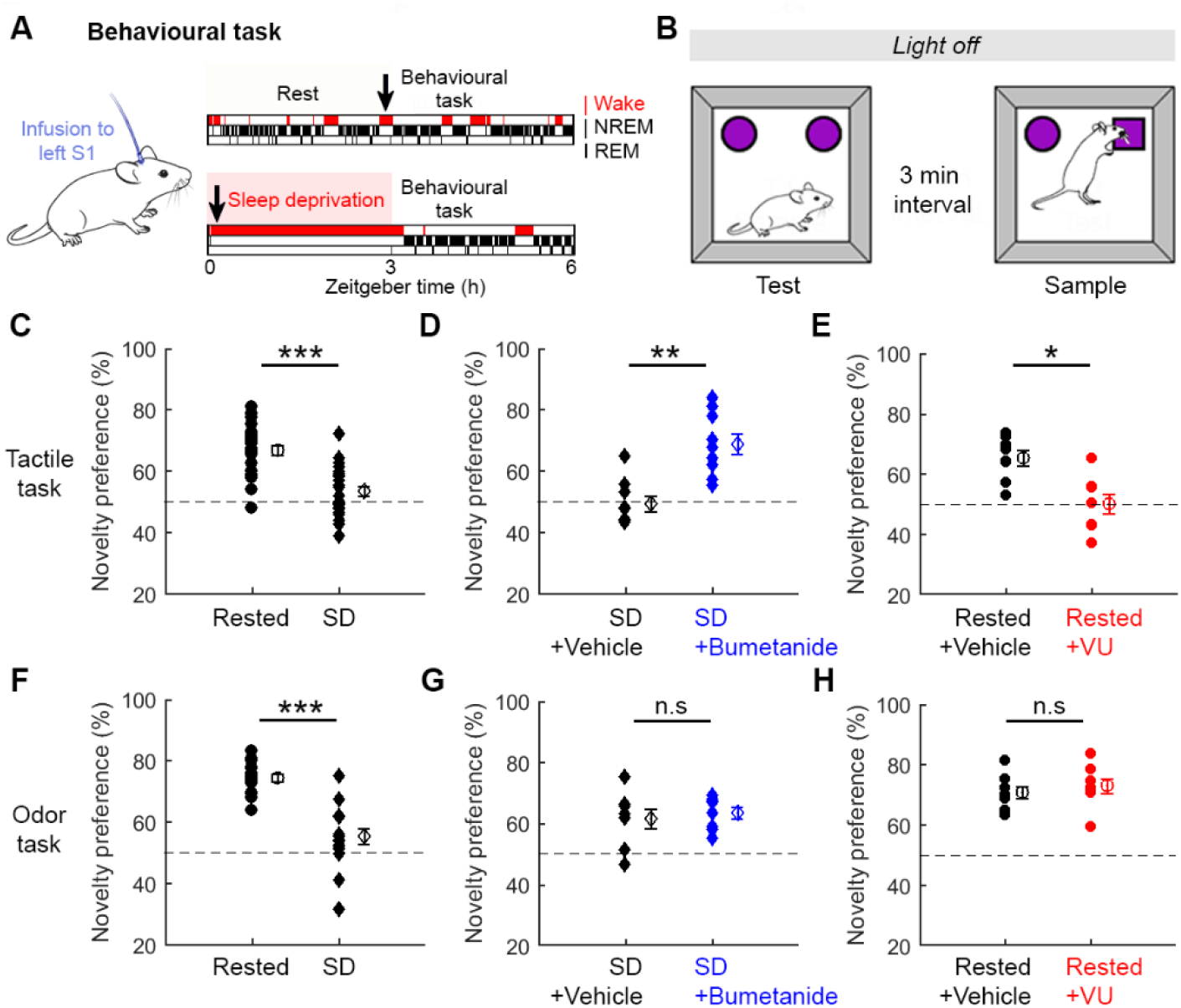
Cortical [Cl^-^]_i_ determines behavioural performance levels in sleep-deprived animals. (**A**) Behavioural performance was tested at ZT3 in either ‘rested’ mice that had been allowed to sleep, or sleep-deprived (‘SD’) mice that had experienced a 3-hour SD protocol at the beginning of the light period. Blockers of NKCC1 or KCC2 were locally infused via a cannula targeted to left S1 (arrows indicate start of infusion). (**B**) A recognition memory paradigm tested novelty preference in either the somatosensory modality (‘tactile task’) or the olfactory modality (‘odor task’). (**C**) SD mice showed lower novelty preference on the tactile task (12 animals, 24 trials; ***p<0.0001 *unpaired t-test*). (**D**) Bumetanide infusion into S1 increased tactile task novelty preference in SD mice (9 animals, 9 trials; **p<0.002 *unpaired t-test*). (**E**) VU infusion into S1 decreased tactile task novelty preference in rested mice (8 animals, 8 trials; *p<0.01 *unpaired t-test*). (**F**) SD mice showed lower novelty preference on the odor task (8 animals, 16 trials; ***p<0.0001 *unpaired t-test*). (**G**) Bumetanide infusion into S1 did not affect odor task novelty preference in SD mice (8 animals, 8 trials; p=0.65 *unpaired t-test*). (**H**) VU infusion into S1 did not affect odor task novelty preference in rested mice (8 animals, 8 trials; p=0.49 *unpaired t-test*).

## Discussion

These data demonstrate that by mimicking the effects of sleep upon local [Cl^-^]_i_ in cortex, it is possible to improve task-specific performance in sleep-deprived animals. Therefore, whilst multiple changes occur with sleep deprivation (*5, 6, 8*), [Cl^-^]_i_ represents a mechanism to overcome these effects in the tired state. This fits with our observations that the [Cl^-^]_i_ of cortical pyramidal neurons changes in a manner that reflects sleep-wake history, is use-dependent, and determines local SWA and low frequency oscillations in the sleep-deprived state. [Cl^-^]_i_ could be considered an activity-dependent ‘tag’, whereby active neurons during waking acquire more depolarized EGABA_A_ and, as a result, a higher probability of being recruited during oscillatory cortical activity that is associated with tiredness.

As Cl^-^ fluxes are principally involved in synaptic inhibition, [Cl^-^]_i_ is uniquely placed to determine how a cortical neuron responds to the synaptic activity patterns within a network, without directly affecting mechanisms that depend upon other ions, such as intrinsic excitability or pre-synaptic release (*28*). The proximity between EGABAA and the resting membrane potential also makes the regulation of [Cl^-^]_i_ attractive from an energetics perspective (*28–30*). Nevertheless, the constant interplay between inhibitory and excitatory synaptic inputs, means that the [Cl^-^]_i_ changes will interact with other effects associated with sleep-wake history. For instance, the high [Cl^-^]_i_ and weakened synaptic inhibition associated with sleep deprivation would be predicted to enhance the recruitment of voltage-gated potassium channels that contribute to oscillatory activity associated with sleep (*31, 32*). As Cl^-^ cotransporters rely upon transmembrane potassium gradients (*33*), there is the potential for cross-talk with neuromodulatory systems that can influence global arousal via extracellular potassium (*7*). [Cl^-^]_i_ also links to evidence that glutamatergic synaptic strength varies with sleep-wake history (*34–36*), as synaptic inhibition gates the induction of glutamatergic synaptic plasticity (*37, 38*), and is therefore relevant to hypotheses that place synaptic plasticity at the heart of the restorative effects of sleep (*40–43*).

Cortical neurons typically display a range of EGABA_A_ values and we identify sleep-wake history and use-dependency as physiological processes that contribute to this variance. Consistent with previous work, the Cl^-^ exporter KCC2 was found to be a major regulator in adult pyramidal neurons (*44*), whilst the contribution of the NKCC1 importer varied as a function of sleep-wake history. The fact that NKCCl’s influence upon [Cl^-^]_i_ was only detected following periods of wakefulness in the mouse, which typically correlates with our night time, may have contributed to this cotransporter being underappreciated in mature cortex. In terms of use-dependency, the potential for Cl^-^ cotransporters to be modulated by activity is well established (*18*). NKCC1 upregulation has been linked to periods of increased activity including in L5 pyramidal neurons (*45, 46*), has been shown to vary on relevant timescales (*47*), and a recent phopshoproteome screen of forebrain identified sleep-wake dynamics in the NKCC1-regulating kinase, WNK1 (*6*). There are also interesting parallels with evidence from the suprachiasmatic nucleus, where diurnal changes in synaptic inhibition have been linked to circadian processes (*47–50*). [Cl^-^]_i_ may represent a fundamental point of convergence between sleep and circadian processes, through which Cl^-^ regulatory mechanisms could translate multiple factors into changes in cognitive performance. Finally, our results provide a new context for understanding neurological disorders such as epilepsy, autism, and schizophrenia, in which altered [Cl^-^]_i_ has been reported in cortical regions, but not yet linked to the sleep disturbances associated with these conditions (*51, 52*). This raises the question of whether altered [Cl^-^]_i_ represents a primary pathology, or a secondary result of changes in sleep-wake patterns. Either way, the co-occurrence of altered [Cl^-^]_i_ and sleep disturbances provides further motivation to target [Cl^-^]_i_ regulatory mechanisms and, by accounting for sleep-wake history, improve therapeutic effects.

## Acknowledgments

We would like to thank the Akerman lab for advice and comments and the Vyazovskiy Lab for support with the in vivo electrophysiology experiments.

## Author contributions

HA and CJA conceptualize the project. HA and CJA designed the in vitro experiments. HA, CJA and VVV designed the in vivo electrophysiology experiments. HA, CJA, VVV, MCP and DMB designed the behavioral experiments. HA performed and analyzed the in vitro electrophysiology experiments. HA and SEN performed the western blot experiments. HA performed and analyzed the in vivo electrophysiology experiments. TY assisted with the optogenetics experiments. HA performed and analyzed the behavioural experiments. PB performed the neuronal network modelling and wrote the automated sleep scoring algorithm. HA and CJA wrote the manuscript with input from all authors.

## Data and materials availability

All data are available in the main text or the supplementary materials.

## Materials and Methods

### Animal husbandry and sleep deprivation

All experiments were performed on male C57BL/6 wild-type mice purchased from Charles River, which were bred, housed and used in accordance with the UK Animals (Scientific Procedures) Act (1986). Animals were maintained under a 12-h:12-h light-dark (LD) cycle. Animals used for sleep deprivation (SD) protocol were singly housed and pre-exposed to novel objects days before experiment to encourage exploratory behaviour. Timing of SD is stated at relevant points in the manuscript (Fig. 1A, 2B, 4A, 5A). The SD protocol then consisted of delivering novel objects under the continuous observation of an experimenter. Once an animal had stopped exploring an object, a new object was presented. This protocol resulted in a mean of 99.2±0.59% (mean ± sem) time spent awake during the SD period.

### Acute brain slices

Acute cortical brain slices were prepared for electrophysiological recordings from 4-12 week old mice at a defined zeitgeber (ZT) time. Time of sacrifice is stated at relevant points in the manuscript (Fig. 1A, 1F, 3A). To prepare acute slices, animals were woken if necessary, and immediately sacrificed by neck dislocation and decapitation. Coronal 350 μm slices were cut using a vibrating microtome (Microm HM650V) in a pre-chilled cutting solution containing (in mM): 65 Sucrose, 85 NaCl, 2.5 KCl, 1.25 NaH_2_PO_4_, 7 MgCl_2_, 0.5 CaCl_2_, 25 NaHCO_3_ and 10 glucose, pH 7.2–7.4 and bubbled with carbogen (95% O_2_/5% CO_2_). Slices were then incubated for at least 1 h in a storage chamber containing artificial cerebrospinal fluid (aCSF; in mM): 130 NaCl, 3.5 KCl, 1.2 NaH_2_PO_4_, 1 MgCl_2_, 1.5 CaCl_2_, 24 NaHCO_3_ and 10 glucose, pH 7.2–7.4, at RT and bubbled with carbogen. When required, slices were transferred to a recording chamber superfused with aCSF, bubbled with carbogen (30°C and perfusion speed of 2 ml/min). Pharmacological manipulations were delivered by bath application of drugs through the perfusion system for at least 10 minutes. Stocks solutions were generated, aliquoted, and stored at −20°C. On an experiment day, stock solution was added to the aCSF to achieve the desired final concentration (in μM): 10 bumetanide (NKCCI inhibitor), 10 VU0463271 (‘VU’; KCC2 inhibitor), 1 TTX (sodium channel blocker), 1 CPG55845 (GABA_B_R antagonist), 3 bicuculline (GABA_A_R antagonist). All drugs were purchased from Tocris Bioscience.

### Gramicidin-perforated patch clamp recordings

To preserve a neuron’s [Cl^-^]_i_ and infer transmembrane gradients for chloride, gramicidin-perforated patch clamp recordings were performed. Patch pipettes were pulled from standard wall borosilicate glass capillaries (2-5 MΩ) and filled with a high KCl based internal solution to be able to monitor the integrity of the perforated patch, and containing (in mM): 135 KCl, 4 Na2ATP, 0.3 Na3GTP, 2 MgCl2, and 10 HEPES. Osmolarity was adjusted to 290 mOsM and the pH was adjusted to 7.35 with KOH. Gramicidin (Calbiochem) was dissolved in dimethylsulfoxide (DMSO) to achieve a stock concentration of 4 mg/ml. This was then diluted into the internal solution on the day of the experiment to achieve a final concentration of 80 μg/ml. The resulting solution was vortexed for 40 s, sonicated for 10 s, then filtered through a 0.45 μm pore cellulose acetate membrane filter (Nalgene) and used immediately.

Neurons were visualized under a 60x water-immersion objective (Olympus BX51WI). Recordings were performed with an Axopatch 1D amplifier (Molecular Devices), acquired using WinWCP Strathclyde software (V.3.9.7; University of Strathclyde) and stored for off-line analysis. Recordings were made when the series resistance had stabilized to ~100 MΩ (approximately 30 min after gigaseal formation). [Cl^-^]_i_ measurements were performed by activating either GABA_A_ or glycine receptors by delivering short ‘puffs’ of either GABA (Tocris Bioscience, 100 μM) or glycine (Tocris Bioscience, 100 μM) via a patch pipette placed in the vicinity of the cell soma and connected to a picospritzer (5–10 psi for 20–40 ms; General Valve). Puffs were delivered at a low frequency (15 s intervals) to ensure recovery of chloride homeostasis. The polarity of GABAergic responses was monitored by activating GABA_A_Rs in current clamp mode at resting membrane potential (I=0). EGABA_A_ measurements were performed in voltage clamp mode from a holding potential of −70 mV. Test voltage ramps (a saw-tooth, down-up function of 500 ms duration, with a minimum of −90 mV and maximum of −50 mV) were delivered at baseline (control ramp) and near the peak of the GABA-evoked current (GABA ramp; Fig. S1A). For each neuron, I-V curves of the control and GABA ramps were generated using linear fits after 70% series resistance correction. EGABA_A_ was defined as the membrane potential at which the control and GABA ramp currents intersected, which was equivalent to the membrane potential at which the difference between the GABA and control ramp currents was equal to zero (Fig. S1B). EGABA_A_ for each data point was a mean of 10 consecutive measurements.

### Western blotting

Animals were sacrificed at ZT3 with or without prior SD protocol. Mouse somatosensory cortex was dissected and the tissue immediately flash-frozen in liquid nitrogen, then stored at −80°C until required. Tissue samples were homogenized in chilled Cell Lysis Buffer (Cell Signalling Technology) supplemented with HALT protease and phosphatase inhibitor cocktail (Thermo Scientific), vortexed, sonicated and incubated on ice for 30 min, followed by 10 min centrifugation at 4°C, 15000 rpm. Total protein levels were quantified with a BCA protein assay (Thermo Scientific) and 40 μg per animal was loaded into a 6% SDS-PAGE gel for electrophoresis. Gels were immunoblotted onto Protran nitrocellulose membranes (Sigma Aldrich), incubated in Intercept (TBS) blocking buffer (LI-COR) for 1 h at RT, and finally incubated overnight at 4°C with primary antibodies against NKCC1 (T4, 1:1000, DSHB) and tubulin (anti-Tuj1, 1:5000, Biolegend) diluted in the blocking buffer with 0.1% tween. The following day, membranes were washed using TBS containing 0.1% tween and incubated with IRDye 680RD secondary antibody (1:5000, LI-COR), diluted in the blocking buffer containing 0.01% SDS and 0.1% tween at RT for 1 h, and shielded from light. Membranes were washed with TBS and imaged with a blot scanner (LI-COR). Each animal’s NKCC1 signal was normalized to its corresponding tubulin signal.

### Surgical procedures and electrode configuration

For chronic electroencephalogram (EEG) and electromyogram (EMG) recordings, custom-made headstages were constructed by connecting three stainless steel screw electrodes (Fine Science Tools) and two stainless steel wires, to an 8-pin surface mount connector (8415-SM, Pinnacle Technology Inc., Kansas). For chronic in vivo LFP recordings combined with drug infusion, custom-designed cannulae (26 gauge) were coupled to two-channel insulated electrode wires (0.125 mm; C315G, PlasticsOne), and connected to the same 8-pin surface mount connector. For 4-channel LFP and multi-unit activity recordings combined with optogenetics, a tetrode in diamond configuration (25 μm spacing) was coupled with a fibre optic (200 μm, 0.22 NA, fibre terminates 100 μm above top site; Q1X1-tet-3mm-121-OCQ4LP, NeuroNexus) and connected to a zif-clip compatible adapter (CQ4-Z32, NeuroNexus).

Device implantation and viral injections were performed using stereotactic surgery, aseptic technique, isoflurane anaesthesia (3-5% for induction and 1-2% for maintenance) and constant body temperature monitoring. Analgesia was provided at the beginning of surgery and during recovery (buprenorphine and meloxicam). A craniotomy was performed over the right frontal cortex (AP +2 mm, ML +2 mm from Bregma), right occipital cortex (AP −3.5 mm, ML +2.5 mm from Bregma), left somatosensory cortex (AP −1.2 mm, ML −3 mm from Bregma) and the cerebellum (~-1.5 mm posterior from Lambda, ML 0). For EEG recordings, a screw was fixed over both the right frontal and occipital cortex. For LFP and multi-unit activity recording, either a cannula coupled with electrodes or a tetrode coupled with fibre optic, was implanted into L5 of left somatosensory cortex (700 nm depth). EEG, LFP and multi-unit activity signals were referenced to the cerebellum screw. For EMG recording, wire electrodes were inserted into the left and right neck muscles and one signal acted as reference to the other. All implants were secured using a non-transparent dental cement (Super-Bond). To express opsins, an AAV1-CamKiia-eNpHR3.0-EYFP or AAV1-CamKii-ArchT-GFP virus was injected, before device implantation, into left somatosensory cortex (40 nl/min, 350 nl at 700 nm depth and 350 nl at 400 nm depth) using a Hamilton syringe (5 μl, 32-gauge) and an infuse/withdrawal pump (Harvard Apparatus). pAAV-CamKiia-eNpHR3.0-EYFP was a gift from Karl Deisseroth (Addgene viral prep #26971-AAV1; http://n2t.net/addgene:26971; RRID:Addgene_26971) while pAAV-CamKiia-ArchT-GFP was a gift from Edward Boyden (Addgene viral prep #99039-AAV1; http://n2t.net/addgene:99039; RRID:Addgene_99039). Animals were allowed to recover for at least 1 week before recordings, or 4-8 weeks to allow for opsin expression.

### In vivo data acquisition

Animals were moved to a recording chamber and housed individually in a Plexiglas cage (20.3 x 32 x 35 cm). Recordings were performed using a 128-channel Neurophysiology Recording System (Tucker-Davis), acquired using the electrophysiological recording software, Synapse (Tucker-Davis), and stored locally for offline analysis (*53*). EEG, EMG, and LFP signals were continuously recorded with a sampling rate of 305 Hz. Extracellular multi-unit activity was recorded at a sampling rate of 25 kHz and filtered between 300 Hz – 5 kHz. Spike thresholds were set manually during the sleep period and used to identify waveform epochs containing spikes (0.48 ms before and 1.34 ms after threshold crossing).

### Vigilance state scoring

LFP, EEG, and EMG data was resampled at 256 Hz and converted into the European Data Format (EDF). The data was pre-processed for automated scoring by computing the spectrograms of the EEG and EMG traces using the multitaper approach (*54*), as implemented in the lspopt module (1 s segments with no overlap, other parameters at default values). Signals in the 0-1 Hz frequency range were removed as these exhibited drift due to animal locomotion, and signals between 45-55 Hz and above 90 Hz were also removed, as these were affected by 50 Hz electrical noise. A log x + 1 transformation was then applied to map the heavy-tailed distribution of power values onto a more normal distribution. Finally, the values in each frequency bin were normalized to a Z-score to ensure that the downstream classification assigned equal weight to all signals across frequencies. To determine vigilance state, we used a hidden Markov model (HMM) to compute the posterior probability of each state based on (i) the probability of each state given the sample, and (ii) the probability of each state given the likelihood of states in previous time points. Both sets of probability distributions were estimated based on training data that had been annotated by human scorers. Linear discriminant analysis (LDA) implemented in scikit-learn (*55*) was used to reduce the dimensionality of the pre-processed and combined spectrograms of the EEG and EMG traces, down to a two-dimensional signal. An HMM with multivariate normal sample distributions for each state and a sparse transition matrix was constructed, and the Viterbi path was computed using the pomegranate module (*56*) to determine the maximum likelihood state sequence. To establish performance levels for the automated method, approximately half of the data sets (55 of the 92 x 24 h recordings from 16 animals) were partitioned into 4 s epochs and manually scored using SleepSign for Animals software (SleepSign, Kissei Comtec) with criteria as described previously (*1*). Automated and manual vigilance state scoring showed a very high correspondence, with 97% match for NREM sleep epochs, 91% match for REM sleep epochs, and 98% match for wake epochs.

### In vivo drug infusion

Stock solutions were made by dissolving drugs in DMSO to reach 10 mM concentration, aliquoting and storing at −20°C. Tubing and internal cannulae were flushed with 70% ethanol and sterile saline the day before the experiment. On the day of the experiment, drugs were prepared by dissolving stock solutions in sterile saline to reach a final concentration of (in μM): 55 Bumetanide and 55 VU0463271. The vehicle contained the same concentration of DMSO in sterile saline. Each infusion solution was loaded into a heavy wall polyethylene tubing (PE50 - C313CT; PlasticsOne) that allowed the animal to move freely, and was connected to an internal cannula (C315I; PlasticsOne) and captive collar (PlasticsOne). A small air bubble was loaded into the tubing, just before connecting to a 1 μl Hamilton syringe (26 gauge needle). In awake mice, the internal cannula was inserted into the implanted cannula at either the beginning of light onset or at the beginning of a SD protocol, by unscrewing the dummy cannula, gently restraining the animal and covering the eyes to evoke brief freezing behaviour. The captive collar was then tightened to secure the internal cannula to the implanted cannula. Infusion was performed using an infuse/withdrawal pump (Harvard apparatus; 700 nl at a speed of 40 nl/min) and monitored by tracking the progression of the air bubble through the tubing. Once the infusion experiment was completed, the internal cannula was disconnected and the dummy cannula replaced.

### Optogenetics activation

Halorhodopsin and Archaerhodopsin were activated by a 561 nm, 50 mW solid-state laser (Cobolt) connected to a 200 μm core patch cord LC-Ferrule (NeuroNexus), which was fixed to the implanted 200 μm fibre optic via a ceramic sleeve. Maximum light power at the fibre optic tip was 30-40 mW and light intensity was further reduced using neutral density filters (ND filters 0.2-0.3; Thorlabs). When mice entered NREM sleep between ZT3 to ZT9, the experimenter activated a protocol consisting of 5 cycles of 20-60s light ON, followed by 40s light OFF.

### Data analysis and statistics

EGABA_A_ measurements were performed using custom Matlab scripts. EEG and LFP spectral power were generated using the Spectrogram function in Matlab, with 0.25 Hz resolution. Spectral power for each frequency bin was normalized to the average power over a 12 h baseline period recorded the day before the experiment, during which the animal was undisturbed. To investigate neuronal recruitment to different phases of a slow wave, the LFP signal during NREM sleep was classified into ON periods (when neurons are more likely to spike) and OFF periods (when most neurons are silent). Detected spikes were plotted on the corresponding Hilbert-transform of the LFP signal (bandpass filtered: 0.5-12 Hz). As recordings were made extracellularly and from deep cortical layers, the OFF period was defined as the upward phase (30 −150° angle) of high amplitude slow-waves waveforms (>30 % LFP mean amplitude), and the remaining phase was classified as the ON period (Fig. S6G-H). These criteria detected a mean of 15.7 ± 2.3 spikes per s during ON periods and 1.87 ± 0.4 spikes per s during OFF periods. All statistical tests were performed using InStat (Graphad). Values within the manuscript are written as mean ± sem.

### Neuronal network simulation

A simple network model was used to explore the effect of EGABA_A_ upon neuronal recruitment during ON and OFF periods. The network was constructed using the neuron simulator Brian 2 (*57*) and comprised 400 glutamatergic neurons and 100 GABAergic neurons. Each neuron was modelled as a single compartment, current-based leaky integrate-and-fire neuron. Free parameters were set as follows: −60 mV resting potential, 200 pF membrane capacitance, 20 ms membrane time constant, 5 ms excitatory post-synaptic potential decay time constant, 10 ms inhibitory post-synaptic potential time constant, and −50 mV spike threshold. The glutamatergic neurons received synaptic connections from the glutamatergic and GABAergic neurons; the GABAergic neurons received connections from the glutamatergic neurons. Within these constraints, the connection probability was set uniformly to 5 %, excitatory synaptic weights were set to 0.3 nS and inhibitory weights were initialized at 0.1 nS. To simulate ON-OFF periods, all neurons in the network received a time-varying excitatory external input consisting of a square wave of a particular frequency (Fig. 3B). The external input comprised two components: the first was common to all neurons and switched between 0 and 100 pA at the defined frequency; the second was a background noise current that was unique to each neuron and varied on a 10 ms timescale, with values drawn from a normal distribution (mean and standard deviation of 50 pA). Before running a simulation, homeostatic inhibitory synaptic plasticity was used to establish a balance of excitatory and inhibitory inputs to each neuron (*58*), and then synaptic weights were frozen. Only EGABA_A_ was then varied and each simulation was repeated 10 times. The full code used to run an individual simulation can be accessed at: https://gist.github.com/paulbrodersen/b6ce791a927b7e7e714e6a322d90e2b7

### Novel object recognition

A recognition memory paradigm was used to test novelty preference within either the somatosensory modality (‘tactile task’) or olfactory modality (‘odor task’). The paradigm was performed in a white rectangular arena (40cm x 60cm x 40cm), under low levels of red light to prevent visual cues. The arena was wiped down with 70% ethanol between sessions. Before the day of the experiment, mice underwent a habituation protocol that involved being placed in the arena for 10 min, on 3 consecutive days. Tactile objects were 3D-printed and varied in shape and surface texture, but were comprised of the same material to avoid odor cues. Exploration time for the tactile objects was defined as the period that the animal’s head was oriented towards the object and the whiskers or paws were in contact with the object. Odor objects such as tea leaves, cinnamon powder or orange peel, were each placed at the bottom of small, open glass jars. This ensured that the odor objects were out of the animal’s reach, to avoid gustatory and tactile cues. Exploration time for the odor objects was defined as the period the animal spent sniffing the top of the jar.

On the day of the experiment, each sample period (i.e. T1) involved placing the animal at one edge of the arena, opposite two identical objects that were each positioned 10 cm away from adjacent corners (Fig. 5B). The animal’s behaviour was then monitored until exploration time reached a total of 60 s or 30 s for the tactile or odor task, respectively. The animal was then returned to its home cage for a 3 min interval, during which the arena was cleaned and a different copy of the familiar object (from T1) was placed in one of the corners, and a novel object was placed in the other corner. The test period (i.e. T2) involved placing the animal at the same starting point in the arena and monitoring behaviour over a period of 5 min. Novelty preference was calculated as the ratio between the time spent exploring the novel object and the total exploration time (Novel/(Novel+Familiar)). The order of vehicle versus drug infusion, and the identity and position of familiar versus novel objects, was each counterbalanced.

---

**Fig S1.**
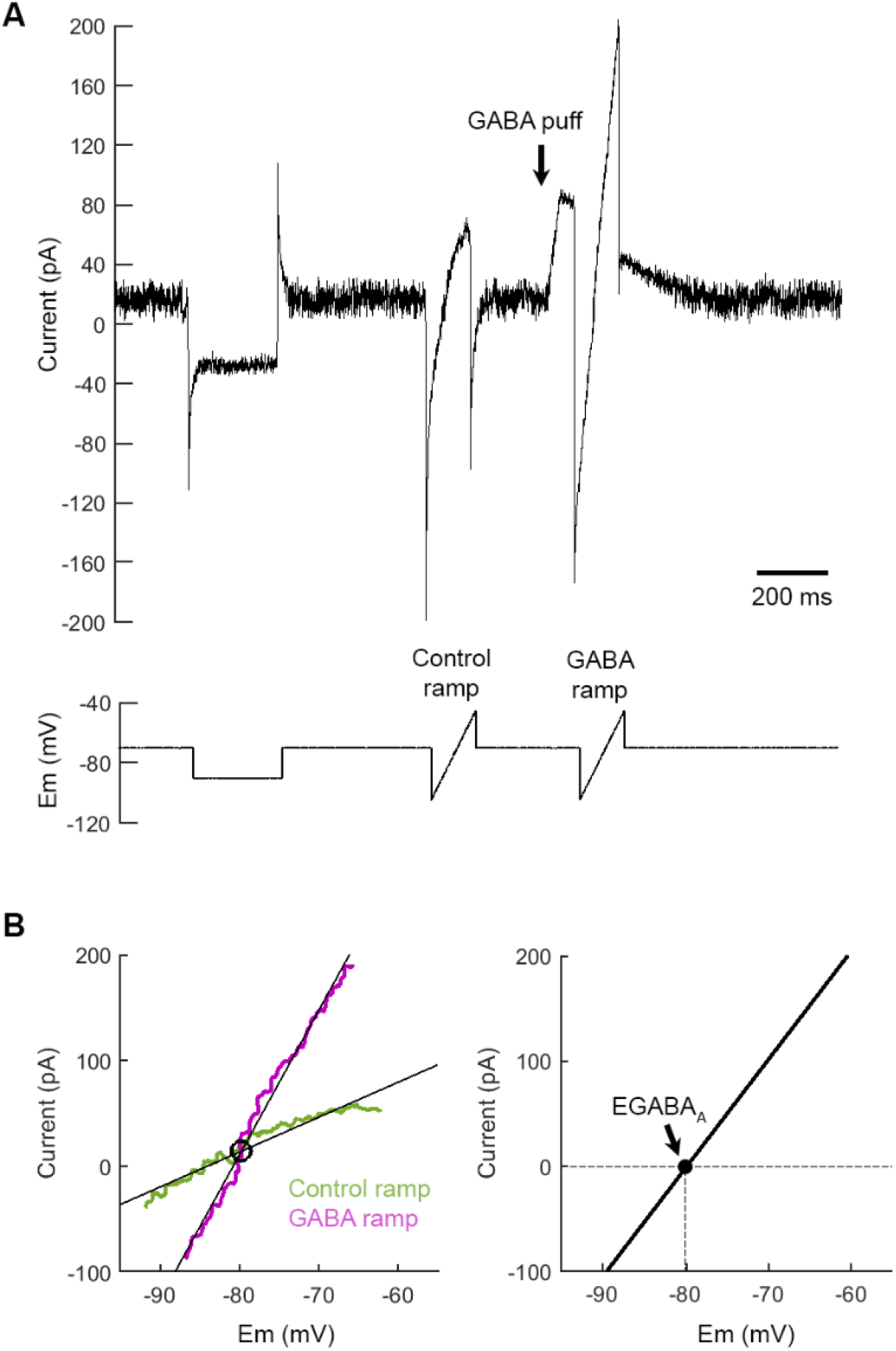
Estimating EGABA_A_ from gramicidin perforated patch recordings. (**A**) S1 L5 pyramidal neurons recorded in voltage clamp mode were subjected to a voltage step and ramp protocol. Voltage steps were used to monitor access resistance. To infer EGABA_A_, a voltage ramp was delivered under baseline conditions (‘Control ramp’) and then at the peak of a GABA_A_R response (‘GABA ramp’), elicited via a GABA puff to the cell soma. (**B**) Current-voltage plots were generated for the control and GABA ramps (left). EGABA_A_ was defined as the membrane potential at which the arithmetic difference between the GABA and control ramp currents was equal to zero (right; black line).

**Fig S2.**
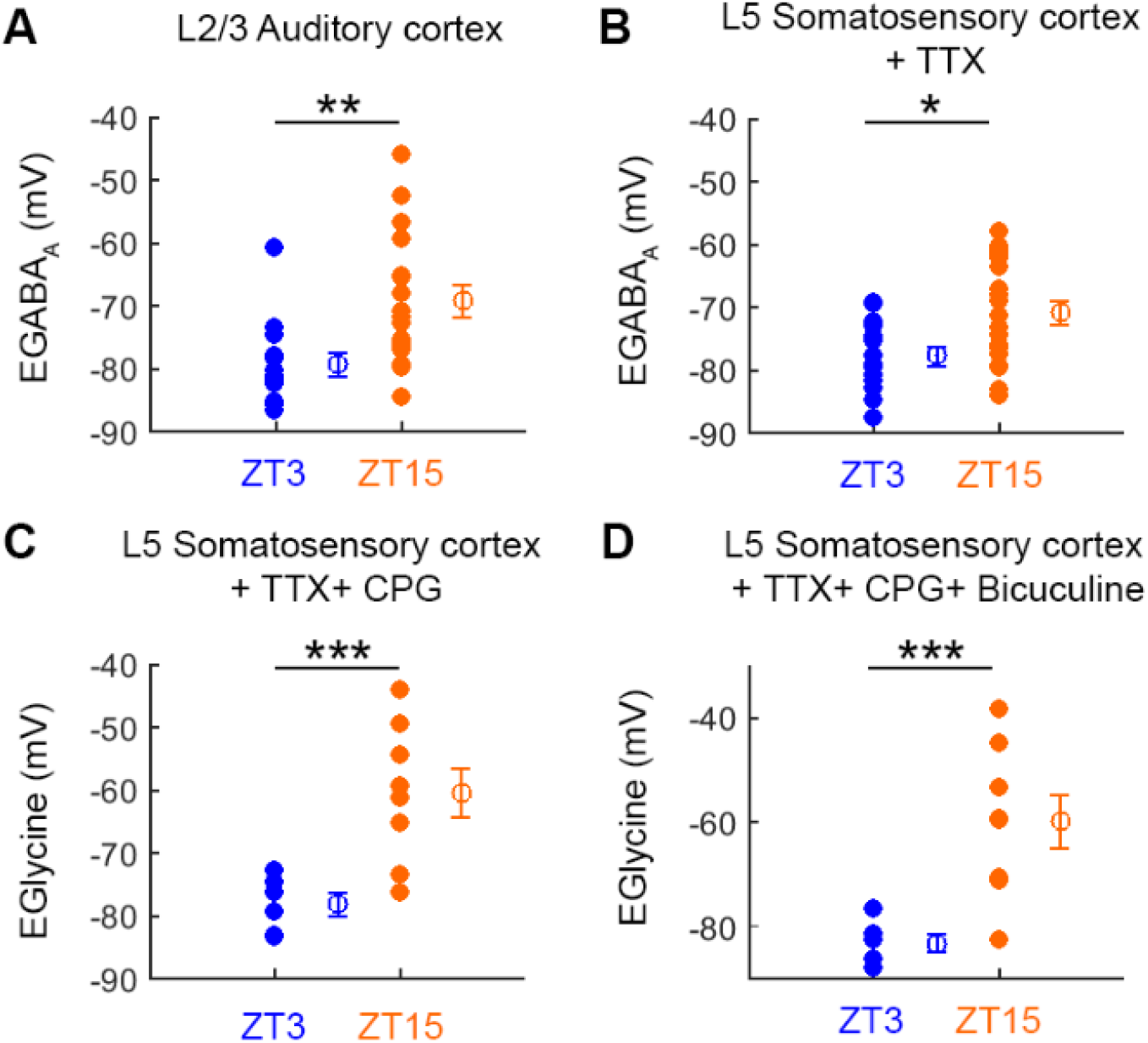
Wake-related changes in [Cl^-^]_i_ occur in different cortical pyramidal neuron populations, and are independent of GABA receptor and spiking activity during the recording. (**A**) In L2/3 pyramidal neurons of auditory cortex, EGABA_A_ was more depolarized at ZT15 (7 animals, 17 neurons) than at ZT3 (7 animals, 13 neurons; **p<0.007, unpaired t-test). (**B**) When network activity in the brain slice was blocked with tetrodotoxin (TTX), the EGABA_A_ of L5 pyramidal neurons in S1 was more depolarized at ZT15 (7 animals, 14 neurons) compared to ZT3 (7 animals, 19 neurons; TTX; *p<0.02, unpaired t-test). (**C**) When GABA_B_ receptors in the brain slice were blocked with CPG55845 (CPG), the reversal potential of chloride-permeable Glycine receptors (EGlycine) in L5 pyramidal neurons from S1 was more depolarized at ZT15 (4 animals, 8 neurons) compared to ZT3 (3 animals, 6 neurons; ***p<0.005, unpaired t-test). (**D**) When GABA_A_ and GABA_B_ receptors were blocked in the brain slice with bicuculline and CPG55845, the EGlycine of L5 pyramidal neurons in S1 was still more depolarized at ZT15 (4 animals, 8 neurons) compared to ZT3 (3 animals, 6 neurons; ***p<0.003, unpaired t-test).

**Fig S3.**
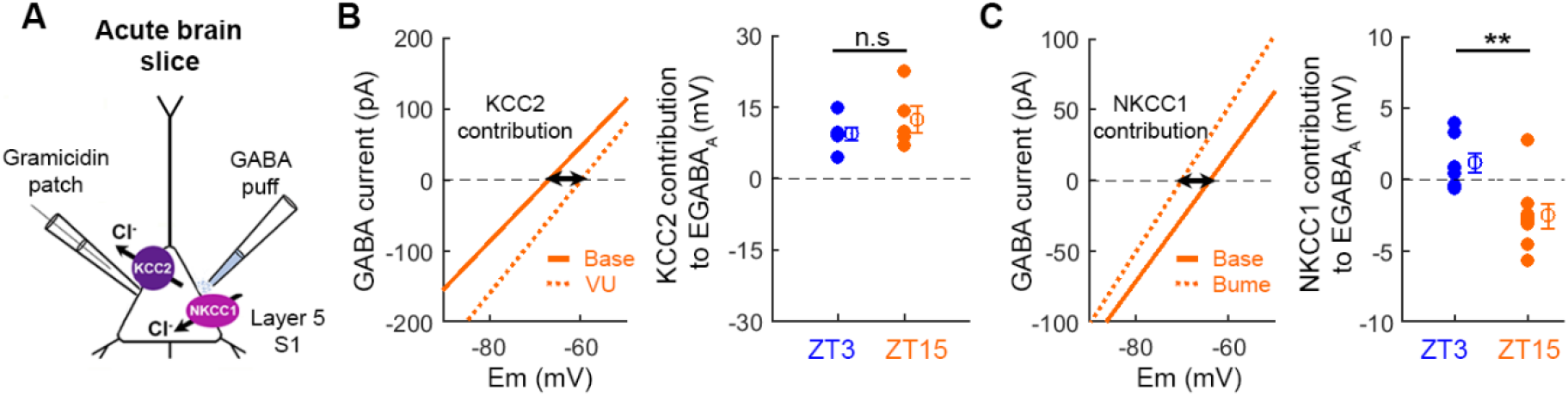
Chloride cotransporter contribution to sleep-wake [Cl^-^]_i_. (**A**) Gramicidin perforated patch recordings compared GABAAR signalling in L5 cortical pyramidal neurons in S1. (**B**) VU0463271 (‘VU’), the KCC2 cotransporter blocker, induced a positive EGABAA shift in a neuron at ZT15 (left). A similar VU-induced positive EGABAA shift was observed in mice at ZT3 (associated with recent sleep; 2 animals, 6 neurons) and mice at ZT15 (associated with recent waking; 2 animals, 5 neurons; p=0.33, unpaired t-test), consistent with a comparable contribution of KCC2 to EGABAA. (**C**) Bumetanide, the NKCC1 cotransporter blocker, induced a negative EGABAA shift in a neuron at ZT15 (left). Bumetanide induced larger EGABAA shifts mice at ZT15 (4 animals, 8 neurons) compared ZT3 (5 animals, 8 neurons; **p<0.003, unpaired t-test). Bumetanide data is repeated from Fig. 1.

**Fig S4.**
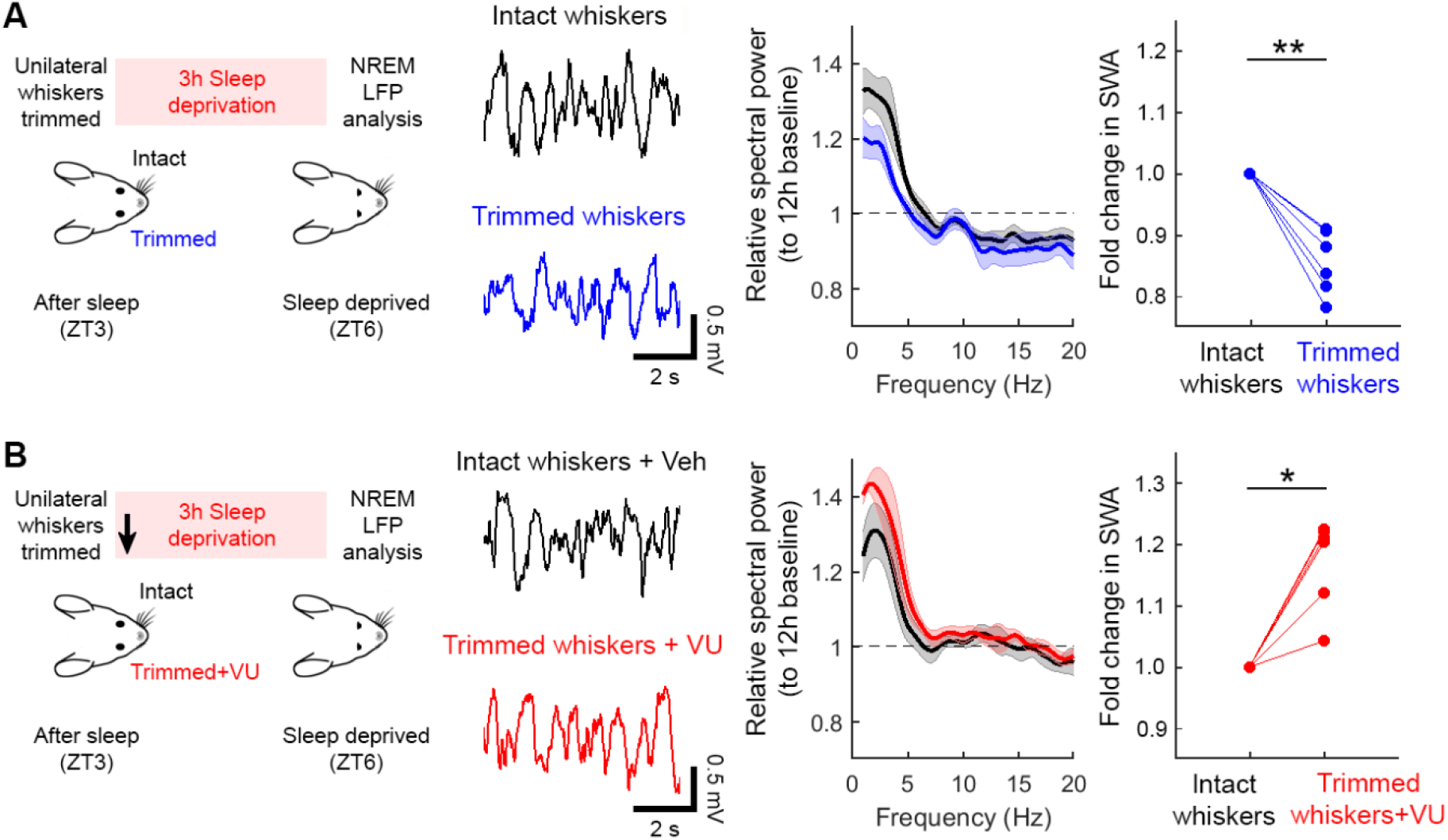
Activity-dependent increases in cortical [Cl^-^]_i_ determine local, use-dependent NREM SWA. (**A**) The same sensory deprivation paradigm was used that had shown wake-related increases in cortical [Cl^-^]_i_ are local and use-dependent (Fig. 1). Whiskers were trimmed unilaterally at ZT3, when [Cl^-^]_i_ is normally low, and mice were subjected to 3 hours of SD. NREM SWA was then analysed during the first 2 hours of sleep following SD. Example LFP signals show SWA recorded from S1 with intact whisker input and from S1 with trimmed whisker input. Data was collected from the same animal, with an interval of at least 2 days. Compared to intact whiskers, trimmed whiskers resulted in reduced SWA during subsequent NREM sleep (right; 6 animals; **p<0.002 paired t-test). (**B**) Using the same sensory deprivation paradigm, [Cl^-^]_i_ was raised in the whisker-deprived S1 by local infusion of the KCC2 blocker, VU (arrow indicates start of infusion). Other conventions as in ‘A’. Pharmacologically raising [Cl^-^]_i_ reversed the reduction in SWA caused by whisker trimming (right; 6 animals, *p<0.003 paired t-test).

**Fig S5.**
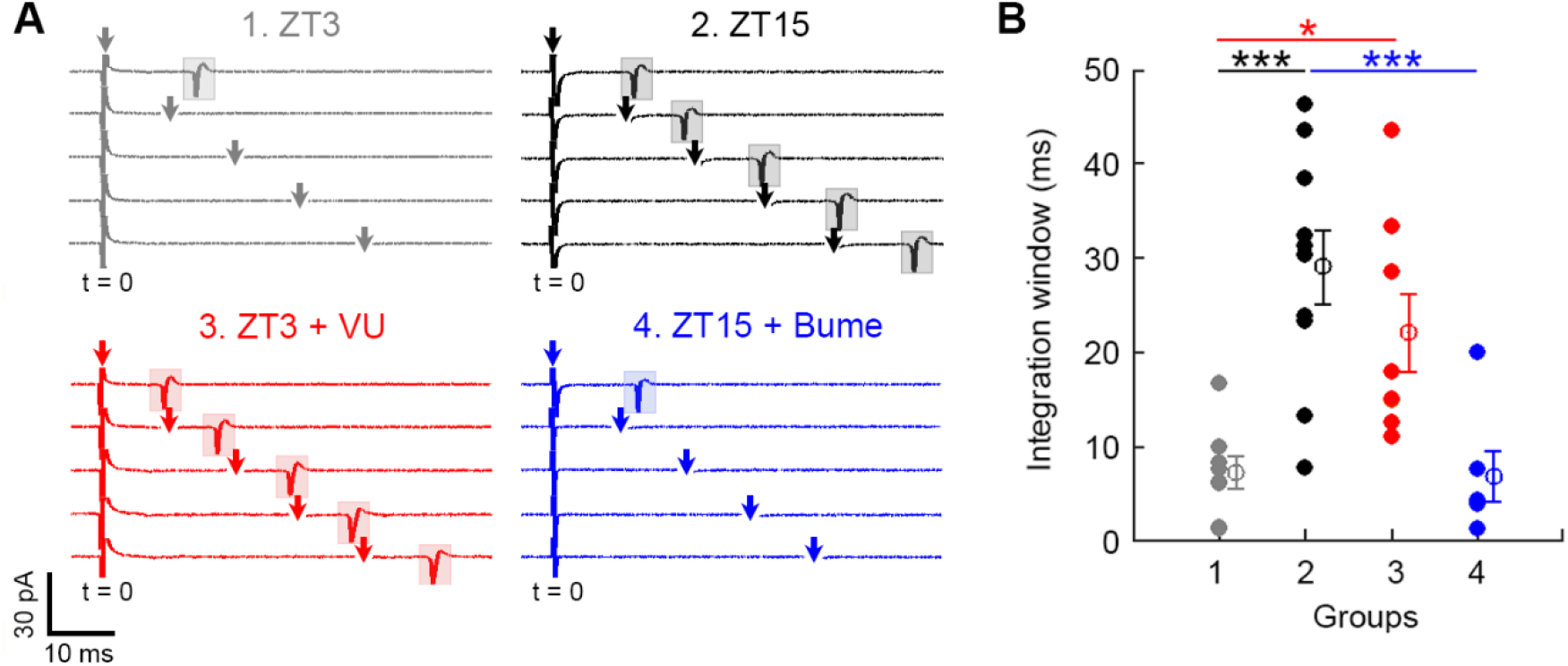
Measuring synaptic integration windows as a function of sleep-wake history and [Cl^-^]_i_. (**A**) Recordings show spiking activity of individual L5 pyramidal neurons in response to the stimulation of two input pathways at different inter-stimulus delays (Fig. 3A). An example is shown for a ZT3 neuron (grey), a ZT15 neuron (black), a ZT3 neuron in the presence of VU (red), and a ZT15 neuron in the presence of bumetanide (blue). The first pathway was stimulated at t = 0, vertical arrows indicate the time that the second pathway was stimulated (artefact removed), and spikes are highlighted by shaded areas. (**B**) Integration windows were defined as the time constant of spike probabilities across inter-stimulus delays (Fig. 3A). Narrow integration windows were associated with ZT3 (grey: 3 animals, 8 neurons) and following NKCC1 blockade at ZT15 (blue: 2 animals, 6 neurons). Wider integration windows were associated with ZT15 (black: 3 animals, 10 neurons) and following KCC2 blockade at ZT3 (red: 2 animals, 8 neurons; multi-group comparison p<0.0001, one-way ANOVA; ZT3 vs. ZT15, ZT15 vs. ZT15 + Bume, and ZT3 vs. ZT3 + VU, all p<0.05, Bonferroni post-tests).

**Fig S6.**
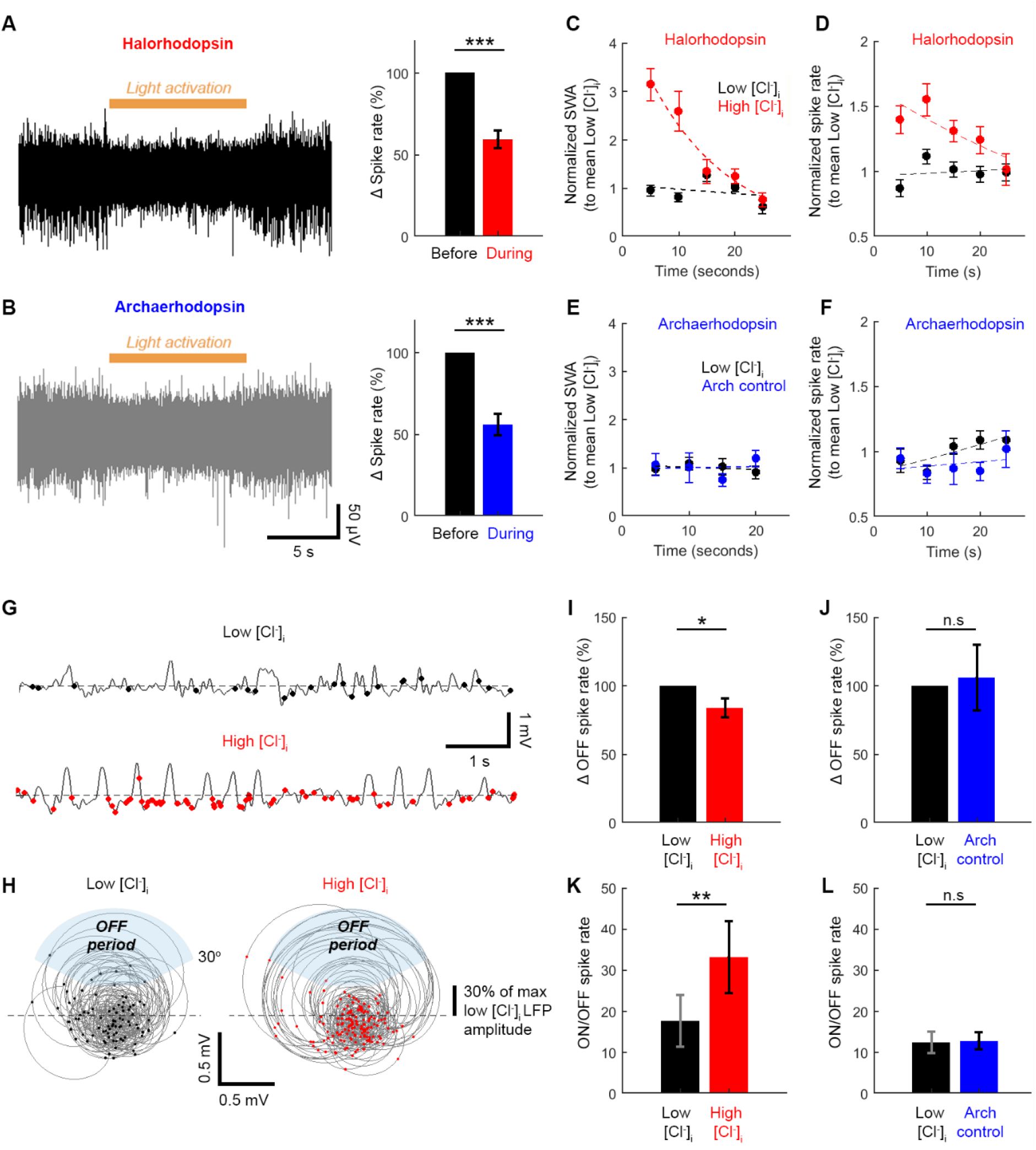
The effects of optical Cl^-^ loading on SWA and neuronal recruitment during ON and OFF periods. (**A**) Representative multi-unit activity from an animal expressing halorhodopsin in pyramidal neurons of S1, during which a 10 s period of light activation was delivered (left). Activation of the opsin was confirmed by comparing spike rate before and during light activation (right). (**B**) Representative multi-unit activity trace from an animal expressing archaerhodopsin (‘Arch’) in pyramidal neurons of S1, during which a 10 s period of light activation was delivered (left). Activation of the opsin was confirmed by comparing spike rate before and during light activation (right). (**C**) Normalized SWA before (low [Cl^-^]_i_) and after (high [Cl^-^]_i_) halorhodopsin activation (5 s time bins) shows recovery kinetics following optical Cl^-^ loading. (**D**) Normalized spike rate before and after halorhodopsin activation (5 s time bins) shows recovery kinetics following optical Cl^-^ loading. (**E**) Normalized SWA before (low [Cl^-^]_i_) and after (‘Arch control’) archaerhodopsin activation (5 s time bins). (**F**) Normalized spike rate before and after archaerhodopsin activation (5 s time bins). (**G**) Representative LFP trace (0.5-12 Hz band pass filtered) during NREM sleep, recorded before (low [Cl^-^]_i_; top) and after (high [Cl^-^]_i_; bottom) halorhodopsin activation. The times of detected spikes are marked with dots. (**H**) Detected spikes were plotted on the Hilbert-transform of the filtered LFP signal. The OFF period was defined as the upward phase (30-150° angle) of high-amplitude SWA waveforms (>30 % of baseline). (**I**) Spike rate during the OFF phase of SWA was reduced after halorhodopsin activation (high [Cl^-^]_i_ condition; 3 animals, 9 days, 22 trials; *p<0.04, paired t-test). (**J**) There was no change in spike rate during the OFF phase of SWA after the Arch control (2 animals, 5 days, 18 trials; p=0.81, paired t-test). (**K**) Normalized spike rate (ON/OFF) during SWA was higher after halorhodopsin activation (high [Cl^-^]_i_ condition; 3 animals, 11 days, 24 trials; **p<0.004, paired t-test). (**L**) There was no change in the normalized spike rate (ON/OFF) during SWA after the Arch control (2 animals, 5 days, 18 trials; p=0.91, paired t-test).

**Fig S7.**
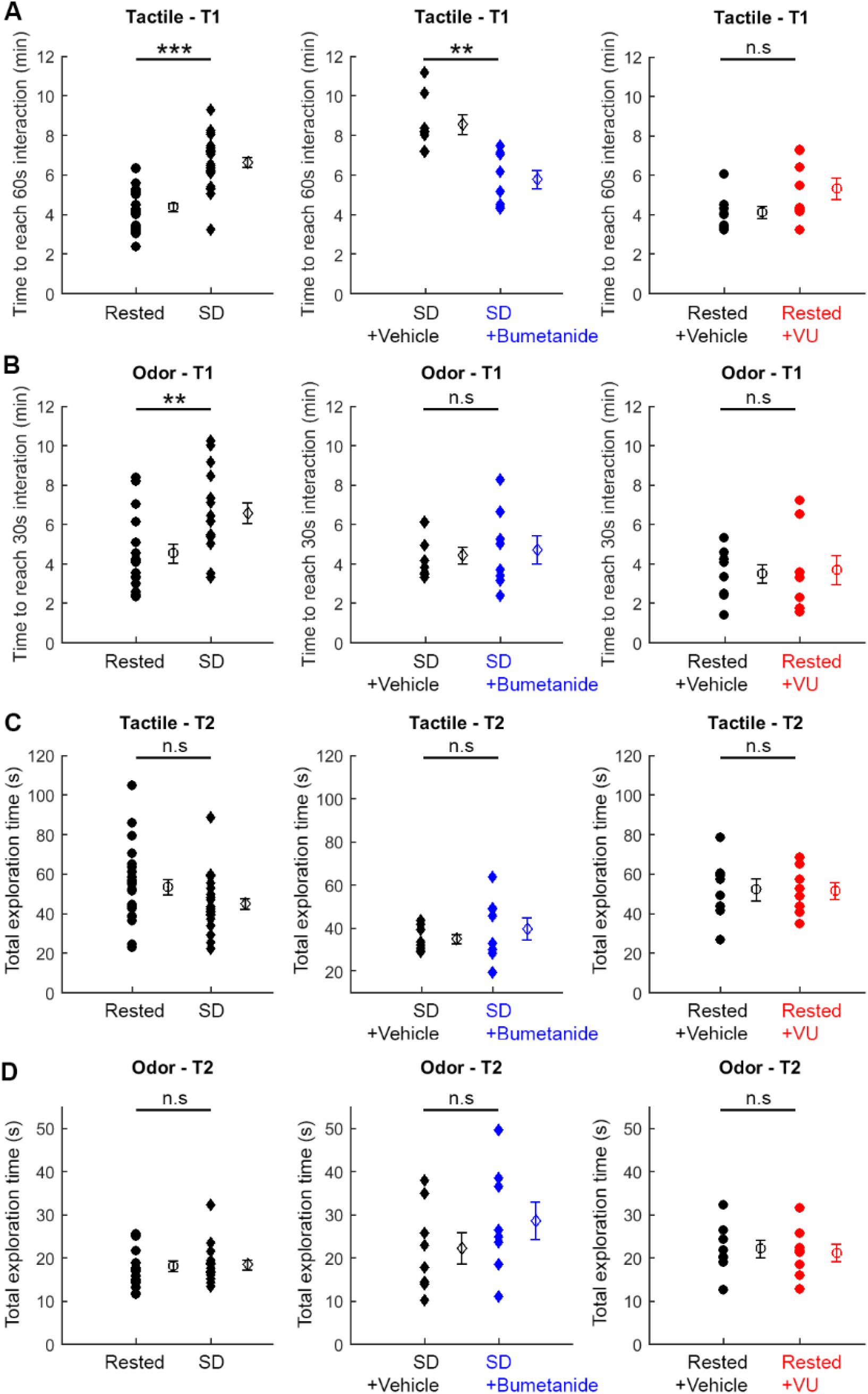
Supporting data from the recognition memory behavioural task. (**A**) Total time before an animal reached 60 s of interaction with the two objects during the sample period (i.e. T1) of the tactile task. Data are shown for rested versus SD animals (left), the effect of a local bumetanide infusion into the somatosensory cortex in SD animals (middle), and the effect of a local VU infusion into the somatosensory cortex in rested animals (right). SD animals took longer to reach 60 s of interaction with the tactile objects (12 animals, 24 trials; ***p<0.0001, unpaired t-test), consistent with a negative effect of SD on performance. SD animals that received bumetanide required less time to reach 60 s of interaction (8 animals, 8 trials; **p<0.002, unpaired t-test). Rested animals that received VU did not show a change in the time to reach 60 s of interaction (8 animals, 8 trials; p=0.08, unpaired t-test). (**B**) Total time before an animal reached 30 s of interaction with the two objects during the sample period (i.e. T1) of the odor task. Plots compare the same experimental groups as in ‘A’. As with the tactile objects in ‘A’, SD animals took longer to reach 30 s of interaction with the odor objects (8 animals, 16 trials; **p<0.007, unpaired t-test), again consistent with a negative effect of SD on performance. SD animals that received a local bumetanide infusion into somatosensory cortex did not show a change in their interaction with odor objects (8 animals, 8 trials; p=0.73, unpaired t-test), supporting a local effect of the NKCC1 blocker. Rested animals that received VU did not show a change in their interaction with odor objects (8 animals, 8 trials; p=0.81, unpaired t-test). (**C**) Total time an animal spent exploring the familiar and novel objects during the first 5 minutes of the test period (i.e. T2) in the tactile task. Plots compare the same experimental groups as in ‘A’. No difference observed in total time exploring tactile objects between SD and rested animals (12 animals, 24 trials; p=0.07, unpaired t-test). SD animals that received bumetanide did not show a change in the time that they explored the tactile objects (8 animals, 8 trials; p=0.4, unpaired t-test). Rested animals that received VU did not show a change in the time that they explored the tactile objects (8 animals, 8 trials; p=0.92, unpaired t-test). (**D**) Total time an animal spent exploring the familiar and novel objects during the first 5 minutes of the test period (i.e. T2) in the odor task. Plots compare the same experimental groups as in ‘A’. SD and rested animals spent comparable amounts of time exploring the odor objects (8 animals, 16 trials; p=0.83, unpaired t-test). SD animals that received bumetanide did not show a change in the time that they explored the odor objects (8 animals, 8 trials; p=0.23, unpaired t-test). Rested animals that received VU did not show a change in the time that they explored the odor objects (8 animals, 8 trials; p=0.76, unpaired t-test).

